# Zinc ions inactivate influenza virus hemagglutinin and prevent receptor binding

**DOI:** 10.1101/2025.04.13.648591

**Authors:** Ahn Young Jeong, Vikram Gopal, Aartjan J.W. te Velthuis

## Abstract

Influenza A viruses (IAV) cause seasonal flu and occasional pandemics. In addition, the potential for the emergence of new strains presents unknown challenges for public health. Face masks and other personal protective equipment (PPE) can act as barriers that prevent the spread of these viruses. Metal ions embedded into PPE have been demonstrated to inactivate respiratory viruses, but the underlying mechanism of inactivation and potential for resistance is presently not well understood. In this study, we found that zinc ions directly impact the binding of influenza virus to host cell sialic acid receptors using hemagglutination assays. Quantifying this effect, we observed that zinc ions inhibit IAV receptor binding within 1 minute of exposure in a concentration-dependent manner. Maximum inhibition was achieved within 1 hour and irreversible for at least 24 hours. Serial passaging of IAV in the presence of zinc did not result in resistance. Overall, these findings are in line with previous observations indicating that zinc-embedded materials impact the IAV hemagglutinin and SARS-CoV-2 spike proteins, and support work toward developing robust, passive, self-cleaning antiviral barriers in PPE.

## Introduction

Emerging infectious agents can have a devastating impact on public health and the global economy (1). Among various emerging infectious agents, RNA viruses are responsible for several recent pandemics and epidemics. Recent and potential future examples include outbreaks of the 2009 pandemic influenza A virus (IAV) strain (2), SARS-CoV-2, the causative agent of COVID-19 (3), and Ebola virus (4). The outbreak frequency of emerging and re-emerging influenza viruses has increased since the 1940s due to intensified human and animal transportation and extensive interactions between humans and the environment (5, 6). While various studies aim to predict or evaluate the risk associated with the next pandemic virus (7–10) and medical advances such as vaccines and antiviral therapeutics can reduce the mortality and symptoms associated with RNA virus infection (11–13), the development of PPE with virus-inactivating properties may help limit the impact of future emerging RNA viruses.

Of particular interest as future pandemic RNA viruses are IAV strains (2, 10). Influenza viruses are the causative agent of flu, an acute respiratory disease with symptoms ranging from a mild fever and sore throat to lethal pneumonia and extra-respiratory complications (5, 14). In humans, flu is primarily caused by IAV and influenza B virus (IBV) strains (15, 16), but some evidence for human influenza C virus infections exists (17). Despite the availability of annual flu vaccinations and antiviral treatments (18), influenza remains a major burden to human health systems, with 3-5 million cases and up to 650,000 deaths reported every year (19). IAVs have also been responsible for four major pandemics since the 1918 pandemic flu (2). The emergence of these pandemic IAV strains, such as the H3N2, H2N2 and H1N1 strains in the last century, is partly driven by an antigenic shift (20–23). This can occur when exchange or reassortment of the hemagglutinin (HA) encoding gene occurs between a seasonal and an avian IAV strain. Concerns about future flu pandemics thus focus on the increased circulation of avian influenza virus strains like H5N1, H5N8, and H7N9, and the ability of these HPAIVs to infect and cause disease in humans (24–28).

The start of an IAV infection depends on the molecular function of the HA protein. The HA protein resides on the outside of IAV particles and binds alpha-2,3 and/or alpha-2,6 sialic acid receptors (29). Following the binding to the sialic acid, IAV particles are internalized into endosomes and the subsequent acidification of these endosomes through viral M2 ion channels triggers mature HA molecules to undergo a conformational change that facilitates fusion of viral and host membranes (30, 31). Membrane fusion starts the release of the viral ribonucleoprotein complexes for transport to the nucleus, where transcription and replication of the IAV genome occurs (32). Antibodies generated upon vaccination or ion channel inhibitors taken orally may prevent different stages of the IAV infection cycle. An alternative strategy to vaccination or drug therapy is inactivation of an IAV particle, for instance through UV radiation, which creates crosslinks in the IAV RNA genome, or virus-inactivating personal protective equipment (PPE) containing divalent ions (33–36).

PPE serve as a first line of defense against infectious pathogens, trapping virus particles, bacteria or fungal spores in a fibrous mesh. This may protect individuals or prevent the spread of a communicable pathogen. Individuals who frequently come into contact with poultry, cattle or wild animals should therefore use PPE and mitigate the risk of human infection and potential outbreaks. We and others have shown that metal ions embedded in nylons or sprays can be used to reduce the titer of viruses and bacteria (33, 34, 37–39). Specifically, we contributed to the development a reusable zinc ion-embedded polyamide 6.6 fiber (PA66) capable of IAV and SARS-CoV-2 (33). However, the mechanism through which IAV and SARS-CoV-2 are inactivated by zinc ions is currently not fully understood and it is unclear if exposure to zinc ions can lead to resistance. A better understanding of the inactivation mechanism is critical to devise better validation assays and further develop self-cleaning PPE.

Here, we demonstrate that exposure of IAV virions to zinc ions impedes sialic acid receptor binding, a critical step for viral entry into host cells. We find that the zinc-mediated inhibition of receptor binding depends on both the concentration of zinc ions and the duration of incubation. The inhibition is retained for 24 hours after zinc ions are chelated with ethylenediaminetetraacetic acid (EDTA), suggesting that zinc ions induce a structural destabilization or conformational change that inactivates IAV HA, thereby preventing cell attachment and viral infection. Six serial passages of IAV in the presence of zinc ions did not result in resistance to zinc inhibition. Overall, our insights provide insight into the mechanism of action of zinc-ion embedded fibers and show that basic virology assays can be used to validate the inactivating properties of self-cleaning PPE.

## Materials and Methods

### Influenza viruses and cells

Madin-Darby canine kidney (MDCK) cells, originally sourced from American Type Culture Collection, were grown in Dulbecco’s Modified Eagle Medium (DMEM) (GeneDepot) supplemented with 10% fetal bovine serum (FBS; Gibco), high glucose, pyruvate and glutamine. MDCK cells were used to amplify influenza A/WSN/33 (H1N1) virus in DMEM with 0.5% FBS at 37°C and 5% CO_2_ or to titrate these viral stocks using plaque assays as described previously (33).

### Zinc incubation and neutralization with EDTA

To assess the effects of zinc or copper ions on IAV, 100 μL of IAV strain A/WSN/33 (H1N1) (7.13 × 10^7^ pfu/ml) was incubated with 100 μL of 0, 1, 2, 5, or 10 mM zinc chloride (ZnCl_2_; Sigma) working solutions prepared in DMEM containing 2% FBS at RT. After incubating the samples for the time periods indicated in the figures, or an hour if not indicated, zinc ions were chelated by adding equimolar EDTA (Sigma), also prepared in DMEM supplemented with 2% FBS, to minimize cytotoxic effects on cell-based receptor binding or infection experiments.

### Hemagglutination assays

To evaluate the effects of zinc ions on sialic acid receptor binding abilities of IAVs, hemagglutination assays were performed using turkey red blood cells (RBCs; Lampire). Two-fold serial dilution of samples were prepared in a U-bottom 96-well microtiter plate (Perkin Elmer) followed by the addition of 1% turkey RBCs in PBS (Gibco) in each well. Plates were incubated for 30 minutes at RT to distinguish between agglutinated and non-agglutinated wells prior imaging. For hemagglutination assays measuring the reversibility of zinc-mediated hemagglutination inhibition, samples were incubated at room temperature for a minute, hour, or a day after zinc exposure and prior to incubation with turkey RBCs. Hemagglutination units (HAU) of samples incubated with zinc ions were normalized to the HAU of each IAV stock used in the absence of zinc ions.

### Passaging assays

Viruses diluted to an MOI of 0.002 for a 6-well plate of MDCK cells were incubated with 2 or 10 mM zinc chloride for 1 h at room temperature. Then zinc ions were chelated by adding equimolar EDTA and the treated viruses added to MDCK cells. After 48 hours, supernatants were collected, frozen and stored as described previously (40). Supernatants were titrated using plaque assay and used to passage the viruses at an MOI of 0.002 6 times. Prior to every infection, diluted viruses were treated with 2 or 10 mM zinc chloride as described above.

### Reverse-transcription PCR

RNA extraction from Trizol was performed as described previously. Spike RNA was purchased from IDT DNA. Isolated RNA was reverse transcribed using SuperScript III and a primer binding to the 3’ end of the RNA. qPCR was performed as described previously. The Spike RNA had the sequence (5’ → 3’): AGUAGAAACAAGGCGGUAGGCGCUGUCCUUUAUCCAGACAACCAUUACCUGUCCACACAAUCUGCCCUUUCGAAAGAUCCCAACGAAAAGAGAGACCACAUGGUCCUUCCUGCUUUUGCU.

### Data analysis and statistics

Data were analyzed in Graphpad Prism 8 using one-way ANOVA with multiple corrections. A Mann-Whitney U test was used to compare the recovery percentages between conditions.

## Results

### Zinc and copper ions reduce IAV titers

Copper and zinc containing surfaces, fabrics and particles have been reported to inactivate IAV strains and SARS-CoV-2. To confirm that copper and zinc ions can inactivate these two RNA viruses in our setup, we incubated a fixed titer of influenza A /WSN/33 (H1N1) with 0-10 mM of zinc or copper chloride. After 60 min, the reactions were stopped with an equimolar amount of EDTA and subsequently diluted for virus titer determination by plaque assay (Fig. 1A). In line with previous observations (33), we found a concentration dependent reduction in both the IAV titer in the presence of zinc and copper chloride (Fig. 1B). In addition, we tested the effect of a mixture of copper and zinc on the IAV and found an increased reduction in virus titer compared to zinc or copper chloride alone (Fig. 1B). We previously showed that EDTA alone does not affect the virus titer (33).

**Figure 1.**
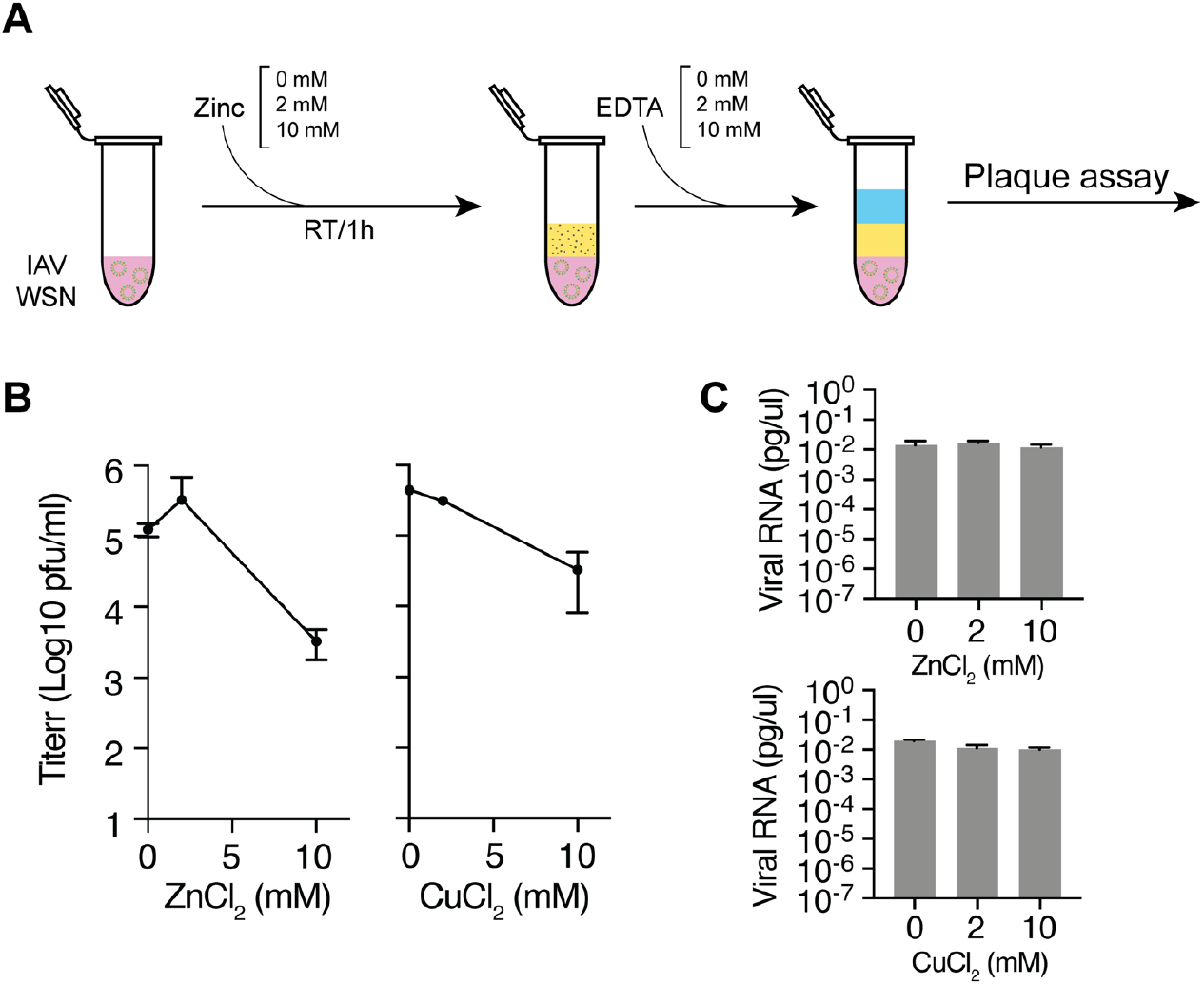
IAV is inactivated by zinc and copper ions. (A) Schematic of the experimental approach for inactivating IAV with zinc or copper ions and neutralization of the zinc or copper ions with EDTA. (B) IAV titers after exposure to zinc or copper chloride and neutralization with EDTA as measured on MDCK cells. (C) NA segment RT-qPCR analysis after exposure of IAV to zinc or copper chloride and neutralization with EDTA.

It was previously shown that IAV and SARS-CoV-2 exposure to zinc ions does not result in viral RNA release (33). To confirm this result, we treated zinc, copper or dually treated IAV with RNase and extracted RNA after a 1-hour incubation. An external non-viral control RNA was added as loading control during the RNA extraction. Next, we performed reverse transcription using an internal or 3’ terminal segment 6 primer and quantified the viral cDNA levels using qPCR. No effect was observed on IAV RNA levels after incubation with zinc or copper chloride (Fig. 1C). Together, these results confirm that zinc and copper ions can inactivate influenza A /WSN/33 (H1N1) in a concentration-dependent manner in our setup and that they do not affect the integrity of the viral particle, in line with our previous observations (33).

### Zinc-mediated inhibition of IAV hemagglutination

Given the relatively low toxicity of zinc, we focused on further characterizing the zinc ions-mediated inactivation of IAV. Building on the above and previous observation that zinc ions leave the viral RNA intact (33), we hypothesized that zinc ions directly affected the IAV HA protein and thereby impaired host cell receptor binding. The gold standard assay for testing HA functionality is the hemagglutination assay, which relies on a change in RBC sedimentation upon the binding of IAV HA proteins to the sialic acid receptors on RBCs. To test whether zinc ions directly affect the interaction between HA and host sialic acid receptors, we measured the sialic acid receptor binding ability of A /WSN/33 (H1N1) after exposure to zinc ions using turkey RBCs.

We first incubated 1 HA unit (HAU) (1.18 × 10^6^ pfu) IAV strain A /WSN/33 (H1N1) with different concentrations of ZnCl_2_ in DMEM supplemented with 2% of fetal bovine serum (FBS). After an hour of incubation at room temperature (Fig. 2A), zinc ions were chelated with an equimolar amount of EDTA to prevent zinc-mediated cytotoxic effects on turkey RBCs during the hemagglutination assay (Fig. 2B). ZnCl_2_ and EDTA-treated IAV samples were next mixed with 1% RBCs and incubated for 30 minutes to allow the RBCs to settle in round-bottom plates. As shown in Fig. 2C, the hemagglutination levels of IAV samples treated with 1 or 2 mM ZnCl_2_ were comparable to RBCs incubated with untreated IAV. RBCs incubated with IAV treated with 5 mM ZnCl_2_ or 10 mM ZnCl_2_ showed a decrease in hemagglutination. The IC_50_ for zinc-mediated inhibition of hemagglutination was 5.00 mM (Fig. 2C). These findings align with our previous results showing a significant reduction in IAV titers after 1 hour of incubation with 10 mM ZnCl_2_. These observations also suggest that zinc ions inhibit IAV HA from binding to sialic acid receptors in a dose-dependent manner, as higher ZnCl_2_ concentrations significantly reduced hemagglutination.

**Figure 2.**
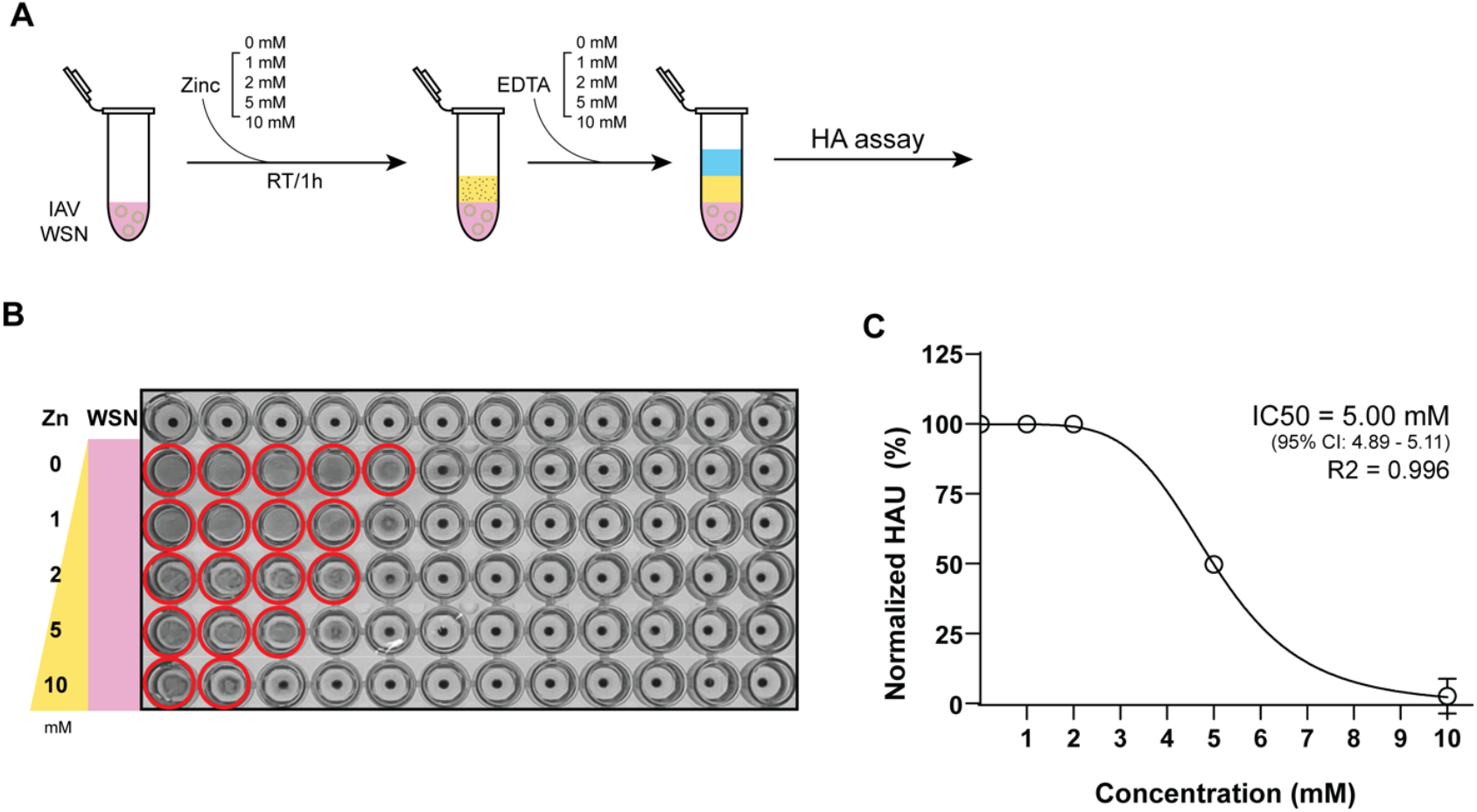
Zinc-mediated inhibition of IAV hemagglutination occurs in a concentration-dependent manner. (A) Experimental design illustrating the evaluation of zinc-mediated IAV hemagglutination inhibition across various zinc ion concentrations and subsequent neutralization with equimolar EDTA. (B) Assessment of sialic acid (SA) binding by IAV virions as measured by the hemagglutination of 1% turkey red blood cells (RBCs) after a 30-minute incubation of IAV with serially diluted zinc ions. Red circles indicate hemagglutination. (C) Hemagglutination units (HAU) of IAV samples treated with zinc ions, normalized to the untreated IAV samples. Nonlinear regression analysis was performed to determine the IC_50_ of zinc-mediated hemagglutination inhibition using a variable slope model. The best-fit IC_50_ was 5.003 mM (95% CI: 4.894-5.111 mM) and the goodness-of-fit analysis yielded an R^2^ value of 0.9960.

### Zinc reduces IAV hemagglutination within 1 minute of incubation

To assess the time required for zinc-mediated hemagglutination inhibition, we examined how different ZnCl_2_ incubation periods affected hemagglutination reduction. Samples were incubated with 10 mM ZnCl_2_ for 1, 10, 30, 60, or 120 minutes (Fig. 3A) and each reaction was terminated using 10 mM EDTA. To minimize variations in ETDA incubation time, zinc exposure of IAV was staggered, with initiation of exposure at different time points and simultaneous termination with 10 mM EDTA. In addition, all hemagglutination assays were performed in parallel, minimizing other confounding factors. Analysis of the hemagglutination level showed that a 1-minute exposure to 10 mM ZnCl_2_ was sufficient to significantly reduce hemagglutination levels compared to the control IAV sample that had been incubated with zinc ions at room temperature for 2 hours (Fig. 3B). The amount of hemagglutination inhibition was comparable among samples exposed to zinc ions for 1 to 30 minutes. Zinc-mediated inhibition was greatest when IAV samples were exposed to 10 mM zinc ions for 1 hour, with no additional reduction observed with longer incubation (Fig. 3C). Using an inhibition curve fit, we calculated the 50% inhibition time to be 1.104 minutes, demonstrating that rapid zinc-mediated HA inactivation occurs within a short time frame (Fig. 3C). These findings indicate that zinc ions exert their antiviral effect on HA rapidly, with significant inhibition occurring within the first minute of exposure and reaching maximal inhibition within an hour, reinforcing the potential for zinc-embedded interventions.

**Figure 3.**
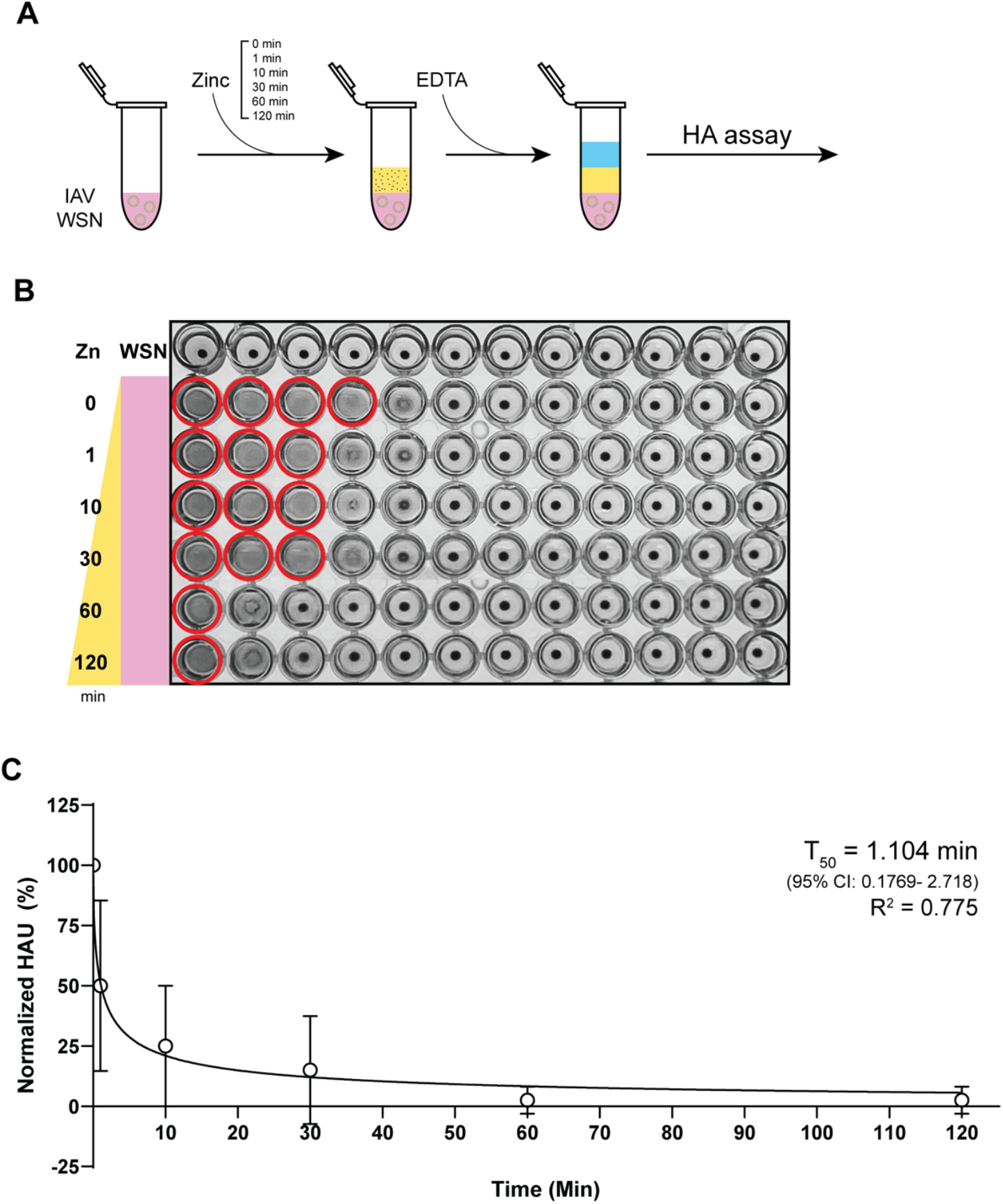
Zinc ions rapidly inhibit IAV hemagglutination within 1 minute of exposure. (A) Experimental design for evaluating the time-dependent inhibition of IAV hemagglutination by zinc ions and subsequent neutralization with 10 mM EDTA. (B) Assessment of sialic acid (SA) binding of IAV virions based on the hemagglutination of 1% turkey RBCs after incubation of the IAV samples with 10 mM zinc chloride for 0-120 min. Zinc ions were neutralized with equimolar EDTA before incubation with turkey RBCs. Solid red circles indicate hemagglutination. (C) Hemagglutination units (HAU) of IAV samples treated with zinc ions, normalized to the untreated IAV control. Nonlinear regression analysis was performed to determine the T_50_ of zinc-mediated hemagglutination inhibition using a variable slope model. The best-fit T_50_ was 1.104 min (95% CI: 0.1769-2.718 mM) and the goodness-of-fit analysis yielded an R^2^ value of 0.7749.

### Zinc-mediated reduction of influenza A virus hemagglutination retained for 24 hours

To assess the durability of zinc-mediated hemagglutination inhibition, we examined how long the antiviral effect persisted after zinc ion chelation with EDTA. After treating IAV with 10 mM ZnCl_2_ for 1 hour and subsequent zinc chelation, the samples were further incubated at room temperature for 1 min, 1 hour, or 24 hours before their hemagglutination levels were assessed (Fig. 4A). Control IAV samples were initially incubated for 1 hour without zinc ions and then for a further 1 min, 1 hour or 24 hours at identical room temperature incubation conditions. As shown in Fig. 4B and C, the hemagglutination inhibition achieved after 1 hour of zinc exposure was maintained for the entire 24-hour period, with comparable levels of inhibition observed. Overall, these findings indicate that zinc-mediated hemagglutination inhibition remains effective and stable for at least 24 hours, suggesting that the antiviral properties of zinc ions persist over an extended period of time, even after zinc ions are removed.

**Figure 4.**
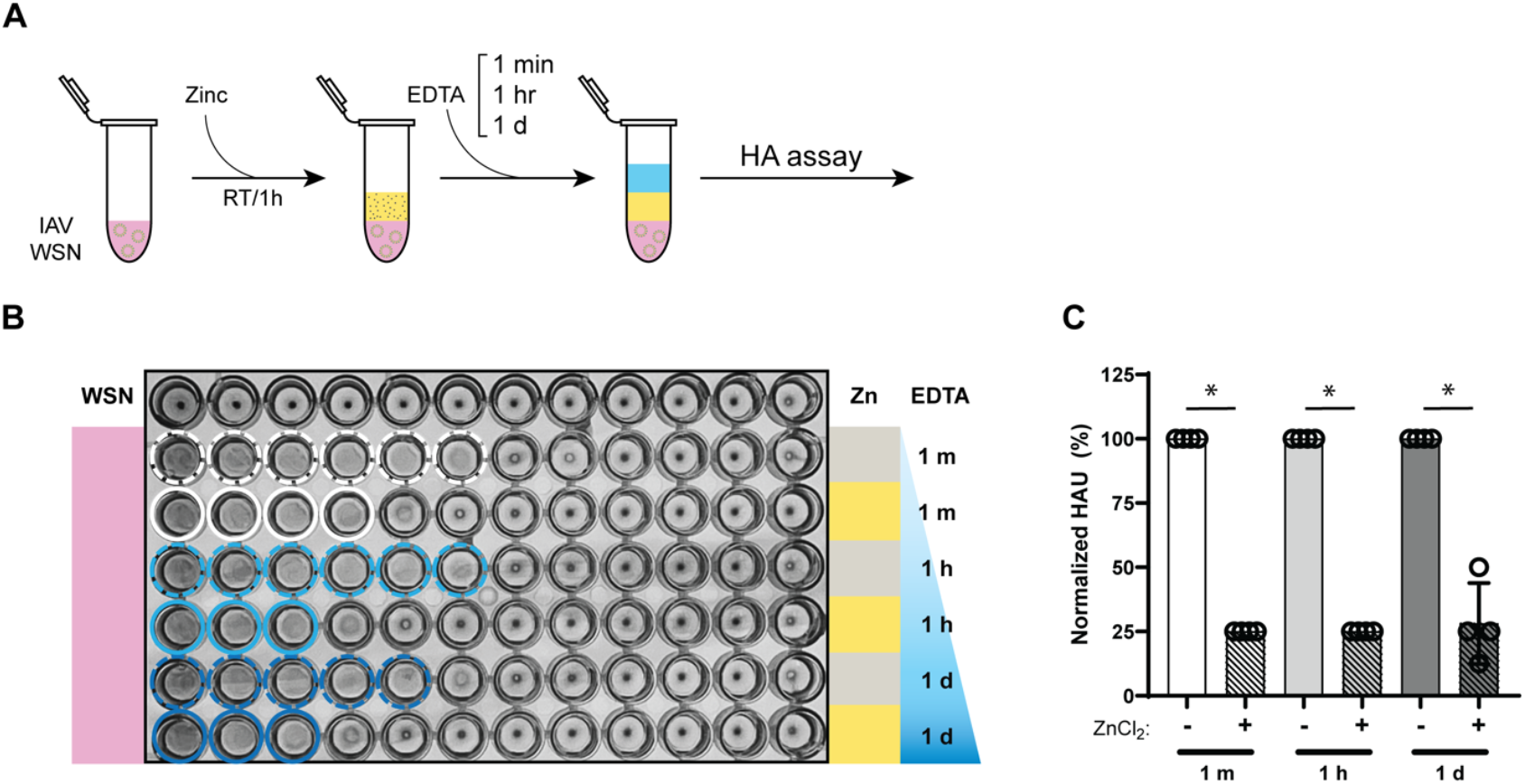
Zinc-mediated HA inactivation is sustained for at least 24 hours. (A) Experimental design for evaluating the persistence of zinc-mediated IAV hemagglutination inhibition for 1 minute, 1 hour, and 24 hours after zinc ion neutralization with equimolar EDTA. (B) Assessment of sialic acid (SA) binding by measuring hemagglutination of 1% turkey RBCs with IAV samples incubated at room temperature for the specified times with zinc (yellow) or no zinc (gray) followed by zinc neutralization with EDTA. Colored solid or dashed circles indicate hemagglutination. (C) Hemagglutination units (HAU) of IAV samples treated with zinc ions, normalized to the untreated IAV samples that were incubated for the same time as the zinc ion treated condition. A Mann-Whitney U test was used to compare the recovery percentages between conditions.

### Zinc-mediated hemagglutination inhibition is maintained across different temperatures

PPE to protect against infectious pathogens may be used in different or fluctuating environments. For instance, face masks experience higher temperatures and higher moisture content than biohazard coveralls (41). Even different parts of a PPE may experience different conditions. The inner surface of a face mask, which is in direct contact with exhaled breath and facial heat, reaches an average temperature of 34.7-35.0°C, while the outer surface is exposed to an average temperature of 34.2-34.9°C. Given that temperature can influence chemical interactions, protein conformations, and ion solubility, we sought to determine whether the antiviral activity of zinc ions against IAV is affected by temperature.

To explore the impact of temperature on IAV inactivation by zinc ions, we incubated IAV with 10 mM ZnCl_2_ at 4°C, 26°C, or 37°C for one hour and chelated the zinc ions with equimolar EDTA at the end of the exposure time (Fig. 5A). The impact of temperature on IAV inactivation was then assessed using hemagglutination assays, which were performed in RT, consistent with our prior experiments. IAV samples incubated at 37°C exhibited zinc-mediated HA inactivation at levels comparable to those incubated at 26°C (Fig. 5B). Samples incubated with zinc at 4°C showed greater variation in hemagglutination inhibition (Fig. 5C), but this variability may stem from reduced solubility of ZnCl_2_ in lower temperatures, as zinc solubility is affected by temperature (42, 43). Overall, zinc-mediated IAV inactivation was significant at both 26°C and 37°C, indicating that the antiviral effects of zinc are robust across a range of physiologically and environmentally relevant temperatures.

**Figure 5.**
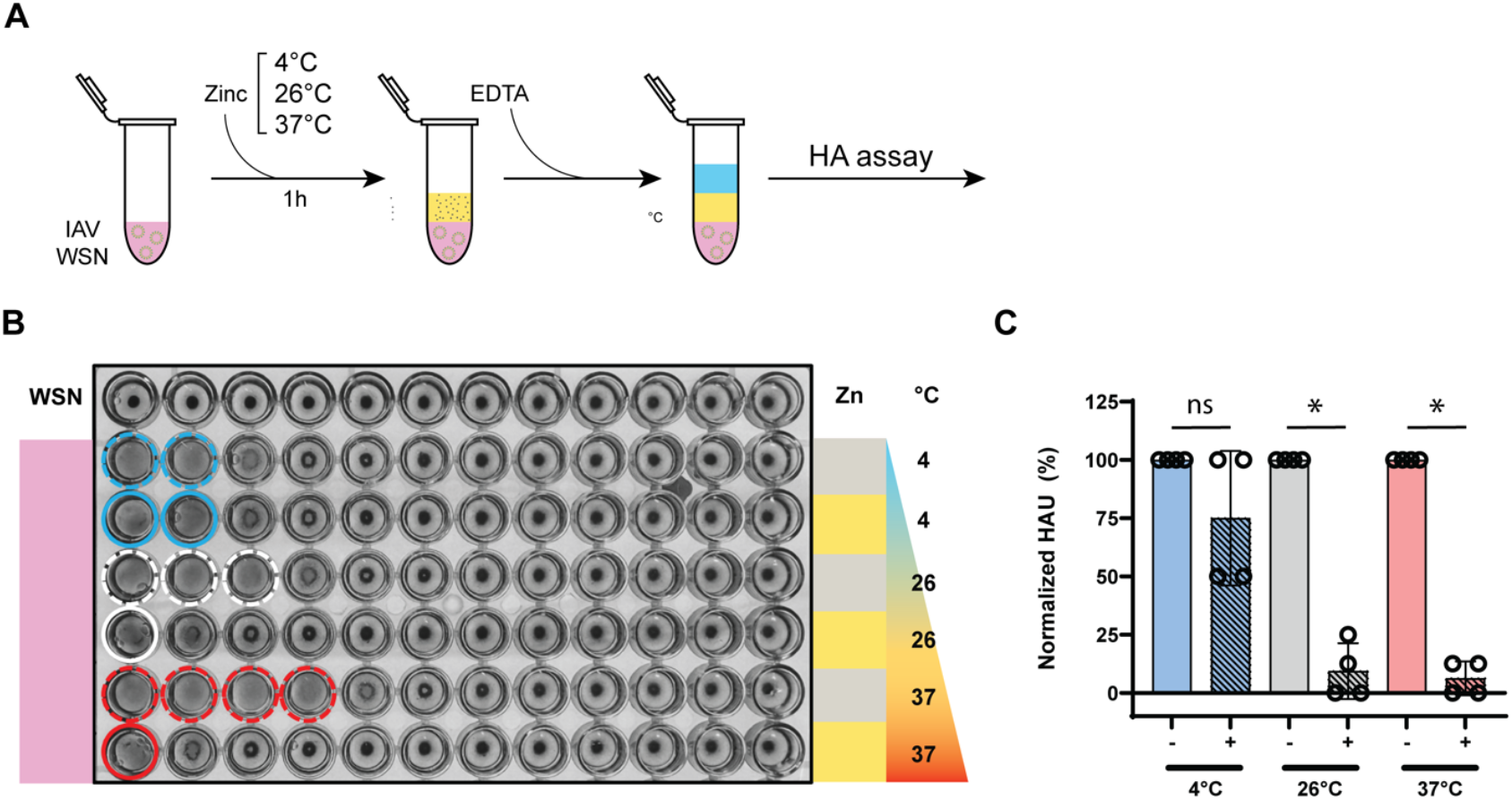
Zinc-mediated HA inactivation remains effective across various temperatures. (A) Experimental design for evaluating the zinc-mediated IAV hemagglutination inhibition at 4°C, 26°C, and 37°C. (B) Assessment of sialic acid (SA) binding by measuring hemagglutination of 1% turkey RBCs with IAV samples incubated at different temperatures in presence (yellow) or absence (gray) of zinc ions for 1 h followed by zinc neutralization with EDTA. Colored solid and dashed circles indicate hemagglutination. (C) Hemagglutination units (HAU) of IAV samples treated with zinc ions, normalized to the untreated IAV samples that were incubated at the same temperature as the zinc ion treated condition. A Mann-Whitney U test was used to compare the recovery percentages between conditions.

### Virus passaging after zinc exposure does not lead to resistance emergence

Exposure of IAV to antivirals, such as favipiravir, can lead to resistance emergence (40, 44). To investigate whether exposure to zinc ions could also lead to resistance, IAV were treated with 2 or 10 mM zinc ions for 1 hour and then used to infect MDCK cells at an MOI of 0.002. After 48 hours, the supernatants were collected, the viruses titrated and the procedure repeated. Overall, we treated and passaged IAV 6 times at the two zinc ion concentrations, but did not observe an increase in titer or decrease in zinc sensitivity (Table 1), suggesting that repeated exposure to zinc ions up to 6 times does not lead to resistance emergence in IAV.

**Table 1.** IAV titers after treatment with zinc ions and 48 h growth (first two columns) or titer after exposure to zinc to measure sensitivity (last two columns).

| Passage | Mean titer after 2 mM zinc passage (pfu/ml) | Titer after 10 mM zinc passage (pfu/ml) | Titer of 2 mM zinc challenge after 10 mM passage (pfu/ml) | Mean titer of 10 mM zinc challenge after 10 mM zinc passage (pfu/ml) |
| --- | --- | --- | --- | --- |
| Untreated | $7.26 \times 10^7$ | $7.26 \times 10^7$ | $1.00 \times 10^5$ (input) | $1.00 \times 10^5$ (input) |
| 1 | $7.83 \times 10^6$ | $8.91 \times 10^5$ | $7.29 \times 10^4$ | $5.13 \times 10^3$ |
| 2 | $6.37 \times 10^7$ | $3.11 \times 10^6$ | $8.98 \times 10^4$ | $4.96 \times 10^3$ |
| 3 | $9.41 \times 10^6$ | $2.83 \times 10^6$ | $4.73 \times 10^4$ | $1.58 \times 10^3$ |
| 4 | $7.17 \times 10^6$ | $6.97 \times 10^5$ | $9.33 \times 10^4$ | $5.73 \times 10^3$ |
| 5 | $4.52 \times 10^7$ | $8.22 \times 10^5$ | $7.11 \times 10^4$ | $8.92 \times 10^2$ |
| 6 | $2.92 \times 10^7$ | $2.09 \times 10^6$ | $5.54 \times 10^4$ | $4.66 \times 10^3$ |

## Discussion

The ongoing challenges posed by emerging RNA viruses, such as SARS-CoV-2 and avian IAV, underscore the need for the continued development of antiviral personal protective equipment (PPE). In particular, there is a growing interest in the application of metal ions, such as silver, copper, and zinc, into nanoparticles and fabrics as a means of providing virucidal activity (45, 46). However, the precise mechanism through which these metal ions exert viral inactivation is still poorly understood.

In this study, we investigated the antiviral properties of zinc ions against IAV. In particular, we focused on their ability to inhibit the host receptor binding function of the IAV HA protein, which mediates viral entry into the host cell. Our findings indicate that zinc ions can inhibit hemagglutination in a concentration-dependent manner (Fig. 2). In addition, our observations indicate that zinc ions can inhibit hemagglutination quickly, with 50% inhibition observed in ∼1 minute of exposure. A maximum inhibition was observed within 1 hour (Fig. 3). In addition, we observed that the antiviral effect of zinc ions was retained for up to 24 hours, indicating a sustained or even irreversible mechanism of action (Fig. 4), and that zinc-mediated IAV inactivation occurred over a wide temperature range, with significant IAV inactivation at 26°C and 37°C (Fig. 5).

Our observations suggest that zinc ions directly affect HA protein function, reducing its ability to bind to host cell receptors. This finding aligns with recent reports that divalent zinc ions induce structural changes in HA similar to those triggered by acidic pH. HA, once primed by proteolytic cleavage from its precursor HA0, undergoes irreversible structural changes in response to the low pH of the endosome, facilitating membrane fusion (47). The precursor HA0 also experiences reversible yet extensive conformational rearrangements at acidic pH, characterized by a wider molecular envelope and dilated and rotated membrane-distal domains. Further structural research is needed to confirm whether zinc ions induce similar conformational rearrangements in HA as a lowering of the pH.

The critical role of PPE in protecting healthcare workers and mitigating SARS-CoV-2 transmission was clearly demonstrated during the COVID-19 pandemic (48–50). However, the pandemic also highlighted significant challenges, such as the risks associated with incorrect PPE use and the incorrect disposal of PPE, which may lead to exposure to virions trapped within the fabric. Our findings, which demonstrate a rapid and sustained inhibition of IAV HA function by zinc ions, highlight the potential of zinc-embedded fabrics as a PPE material with immediate virucidal activity that is long-lasting and active across a range of temperatures. By preventing virus entry into host cells, this fabric could significantly reduce the risk of infection at its earliest stage, which would be of value in high-risk environments, such as healthcare settings. Moreover, the sustained protection provided by zinc-embedded PPE would not only stop the virus from initiating replication, but also improve safety during and after use.

Overall, these findings reinforce the potential utility of zinc-based antiviral strategies in PPE applications, particularly in settings where temperature fluctuations occur. The observed consistency in zinc-mediated HA inactivation at higher temperatures supports the feasibility of zinc ion-embedded materials as self-cleaning antiviral barriers for face masks and other protective equipment. With avian IAV spillovers becoming more frequent, especially among individuals who are in close contact with livestock, effective virus-inactivating PPE could be a key measure in reducing the occurrence of zoonotic transmission and preventing the next pandemic. While we here based our work on the A/WSN/33 (H1N1) strain and additional studies are required to confirm that zinc can offer similar protection efficiencies against avian influenza strains, we think that our work advances our understanding of the virucidal properties of zinc and hope that it will inspire further research.

